# Sensorimotor dynamics of target acquisition and homing in human echolocation

**DOI:** 10.64898/2026.06.13.732093

**Authors:** Santani Teng, Giovanni Fusco, Aarshil Patel

## Abstract

Blindness imposes constraints on the acquisition of environmental sensory information. To mitigate those constraints, some blind people employ *active echolocation*, a technique in which self-generated tongue “clicks” produce informative reflections from surrounding surfaces. Practitioners typically produce multiple clicks that guide, and are in turn shaped by, goal-relevant action. What perceptual information is gained in the echoacoustic signal from each click, and how does it inform motor behavior during task performance? To explore these poorly understood dynamics, here we recorded head movements and clicking behavior of an early-blind expert echolocation practitioner who localized and oriented toward a target object positioned at a 1 m distance and random azimuth in the frontal hemifield. Three additional participants, including a blind self-reported echolocator, were unable to perform the task better than chance level. Performance clearly benefited from available echoacoustic information: The larger target was localized with an average absolute angular error of 9.5° in 9.3 s, vs. 24.6° in 23.2 s for the smaller target. In a passive control condition prohibiting clicks altogether, no significant convergence on the target was observed, confirming the necessity of active sampling. Clicks were emitted somewhat more rapidly and intensely for small targets, but within-trial emission rate and head kinematics (left-to-right reversals) remained relatively invariant. Angular convergence toward the target was consistent with an exponential decay profile, though only weakly distinguishable from a linear trend for small targets. Pooled across trials within each condition, clicks were unimodally distributed about the target azimuth, suggesting an intensity-maximization strategy. In sum, clicking behavior and target size (therefore sonar strength) strongly influenced the rate and precision of orientation convergence toward the target, suggesting that dynamic interactions between motor-driven head movements, click production, and the resulting echoacoustic feedback accumulate goal-relevant evidence across multiple samples. Together, these results illustrate naturalistic sensorimotor dependencies underlying auditory active sensing in the absence of vision.

## INTRODUCTION

In the absence or insufficiency of vision, the critical function of perceiving and interacting with the environment is mediated by nonvisual sensory modalities. Like bats and dolphins, some blind humans address this challenge using *active echolocation*: they ensonify their surroundings with a self-generated sound and extract object and spatial information from the resulting echoes. That sensory signal, in turn, drives goal-directed motor behavior such as locomotion and/or further click production. While ensonification methods vary (Kellogg 1962; Rojas et al. 2009, 2010; Schenkman and Jansson 1986; Bujacz et al. 2022), the typical self-generated signal in studies of echolocation is a sharp palatal or alveolar oral vacuum pulse, or “click,” produced with the tongue (Thaler et al. 2017). This technique enables echoacoustic spatial perception (Teng et al. 2012; Thaler et al. 2019; Rice et al. 1965; Thaler et al. 2022; Flanagin et al. 2017; Teng and Whitney 2011), object and material discrimination (Milne et al. 2014; Teng et al. 2024; Arnott et al. 2013; Hausfeld et al. 1982), and navigation (Rosenblum et al. 2000; Fiehler et al. 2015; Dodsworth et al. 2020; Thaler et al. 2020).

Even beyond navigation, echolocation depends critically on motor actions, including self-motion and the click production itself. Echoacoustic spatial and object judgments are facilitated by head movements (Tonelli et al. 2018; Milne et al. 2014; Wallmeier and Wiegrebe 2014), walking (Rosenblum et al. 2000), and repeated click emissions (García-Lázaro and Teng 2026; Thaler et al. 2018). Echolocators tend to make fairly stereotyped clicks — they are roughly broadband, differing across individuals but highly consistent within individuals (Thaler et al. 2017).

Observers may make more or louder clicks in response to more challenging conditions (Thaler et al. 2018, 2019) but generally have not been reported to alter click timing or other click acoustics. In lab-controlled psychoacoustic settings, more repetitions of identical synthetic stimuli systematically improve localization thresholds (García-Lázaro and Teng 2026). These results are interpreted as supporting a dynamic evidence-accumulation process in echolocation, where multiple sensory samples are integrated toward a percept. However, real-world echolocation happens in concert with goal-directed motion, and the sensorimotor dynamics of localization behavior have only rarely been tested behaviorally. Separately, the extraction of spatial information from echoes is not simply a function of click acoustics, but depends on observer-controlled choices. For example, free-moving Egyptian fruit bats (among the few “clicking” vs. “chirping” species) were found to ensonify their targets bimodally — slightly off-axis to either side such that the target fell within the region of steepest slope, rather than maximum intensity, of the click energy distribution (Yovel et al. 2010). Human clicks are far less directional, essentially isotropic within a roughly 60° cone (Thaler et al. 2017). Interestingly, human echolocalization thresholds are also finer 45° off-axis compared to the midline (Thaler et al. 2022), an effect attributed not to the click energy distribution but to steeper echo-specific binaural cue slopes at off-axis eccentricities. Still, it has not been tested whether human echolocators maximize echoacoustic intensity or slope under realistic (e.g. head-free) behavioral conditions. Thus, the dynamic, online processes by which motor actions and echoacoustic perception combine to meet behavioral goals remain only sparsely understood.

Characterizing human echolocation in an active-sensing framework, including ecological motor behaviors such as self-motion, would place it in the context of other active-sensing behavior, such as humans’ visual exploration of a scene via sequential eye movements (Yang et al. 2016, 2018; Renninger et al. 2007), and dynamic echolocation behavior in nonhuman species such as bats (Yovel et al. 2010; Ulanovsky and Moss 2008; Moss and Surlykke 2010; Macias et al. 2022). Such an approach would clarify the perceptual mechanisms underlying echolocation, and at a practical level, could serve as a basis for optimizing training regimens, assistive technology, or other therapeutic interventions.

To this end, here we measured the time course of motor and sampling behavior (head movements and click emissions), the associated sensory input (auditory echo returns), and the resultant performance (azimuthal response accuracy and precision) as an experienced blind echolocation practitioner performed a target acquisition and localization task. Three additional participants attempted the task but did not show systematic convergence (see Methods & Appendices). Thus our detailed single-case report characterizes an established active-sensing process reflecting a high, stable level of echolocation proficiency. We show that both accuracy as well as target acquisition (head movement) behavior are modulated by the size of the target and the availability of actively generated click information, and that human echo-acquisition strategy likely capitalizes on different acoustic cues from that of clicking bats.

## METHODS

### Participants

Four participants, two blind and two sighted, were initially enrolled in the experiment (Table 1). All participants provided informed consent in accordance with protocols approved by the Smith-Kettlewell Institutional Review Board. Initial testing revealed that three of the four were unable to perform the task at better than chance levels nor to complete every condition in the experiment (see supplemental data in Appendix 1). This included an additional late-blind participant, with self-reported regular use of echolocation since becoming blind in adolescence. Thus here we focus on an early-blind (“EB”) male, highly proficient daily echolocation practitioner, age 53 y, completely blind since infancy due to surgical enucleation.

**Table 1.** Participant characteristics.

| Subject | Sex (M/F) | Visual Status | Age of blindness onset (y) | Age at test (y) | Self-reported echolocation use |
| --- | --- | --- | --- | --- | --- |
| SB01 (EB) | M | Blind (total) | 0 | 53 | Daily |
| SB02 | M | Blind (total) | 16 | 70 | Daily (indoor) |
| SS01 | M | Sighted | N/A | 17 | None |
| SS02 | F | Sighted | N/A | 40 | None |

### Apparatus and stimuli

The experiment was conducted in a soundproof, double-walled sound- and echo-attenuating chamber (IAC Acoustics, North Aurora, IL). A ceiling-mounted rod turned on an axis directly above the participant’s chair, controlled manually by the experimenter (Figure 1A). At a radial distance of 1 m, a second thin rod extended down to the participant’s head level to present the target — an aluminum-foil-covered foamboard rectangle, 1 cm thick and 29 cm wide x 36 cm tall. Used for the “Big Target” condition, it subtended 17° azimuthally from the rod’s center of rotation, where the participant sat. In the more difficult “Small Target” condition, the rod held a much smaller target — a strip 2.5 cm wide x 17 cm tall, subtending 1.4° azimuthally (Figure 1B).

**Figure 1.**
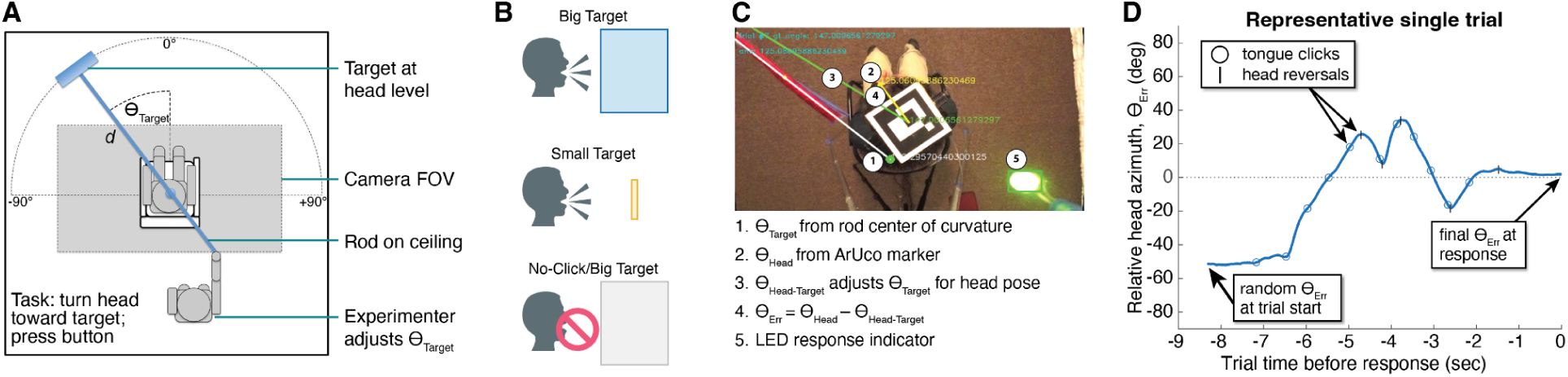
Experimental setup, task, and basic analysis. **A**. Schematic overview of setup within testing chamber. Radial rod mounted above the participant; target at head level. **B**. Experimental conditions: echolocalizing a large (*top*) or small (*middle*) target, or attempting to do so without echolocation (*bottom*). **C**. Example video frame, showing extraction of θ_Err_ and response indicator from extracted image features. **D**. Single-trial time course of angular error *θ*_Err_ between head and target, counting down to response at t = 0.

### Task and experimental procedure

Each trial began with the participant pointed straight ahead (0°) with ears covered. The experimenter positioned the reflecting target at a pre-randomized range of azimuths in the frontal hemifield, up to roughly ±100° relative to EB’s heading. Upon trial initiation via shoulder tap from the experimenter, EB began the task of detecting and localizing the reflector as accurately as possible, following the instruction to “point your nose at the center of the target” using echolocation and then to end the trial via a button press. The head-pointing paradigm is ecological to the task and well-established in auditory localization studies (Pralong and Carlile 1996; Makous and Middlebrooks 1990; Zwiers et al. 2001). No time limit was imposed. This procedure was the same for both Big Target and Small Target conditions. Additionally, we included “No Click” control trials in which a large target was present, but EB had to attempt the task without making any active clicks — to determine, e.g., if incidental apparatus noise, experimenter movements, or ambient sound were informative to the task.

To balance experimental control with ecological validity, we did not fix EB’s head beyond instructing him to remain seated in the chair. This meant the head was not always exactly coaxial with the apparatus (contrast with Rice 1969), but we tracked head pose relative to the target via overhead video for post-hoc analysis (see next section). We also accommodated the greater physical strain of localizing targets in the Small Target condition, where many more clicks were required per trial, and the “catch trial” function of the No-Click condition. Thus, in total, EB performed 103 Big Target trials, 50 Small Target trials, and 28 No-Click control trials over three sessions spanning two days, with nonparametric analyses accounting for the unequal sample sizes (see below).

### Data acquisition and processing

Sessions were recorded from overhead with a video camera (Hero5 Session, GoPro Inc.) at 60 Hz and 1920×1080 resolution, with audio recorded at 48 kHz to mark echolocation click events.

Additionally, we recorded audio via binaural in-ear microphones and a digital recorder (SP-TFB-2 microphones, Sound Professionals, Penndel, PA; DR-2D recorder, Tascam, Tokyo, Japan) at 96 kHz and 24 bits. The concha-seated microphones did not occlude the observer’s ear canals, and allowed us to capture subject-centered echoacoustics for further analysis.

#### Head and Target Tracking

To monitor head pose over time, we fitted the participant with a cap displaying an ArUco marker, a binary matrix image used for fast detection and pose estimation within the OpenCV framework (Garrido-Jurado et al. 2014; Bradski and Kaehler 2000) (https://opencv.org). Additionally, the rod holding the echo reflector was affixed with a direction indicator angled to always be within the camera’s field of view (Figure 1C). To minimize distortions arising from misalignments between camera position and rod center of rotation (CoR), we estimated the CoR empirically for each block. Finally, a green LED illuminated a portion of the camera view whenever the subject responded via button press, which allowed us to segment trials by their final video frame. Thus, after calibrating for camera distortions and measuring constant physical dimensions, we extracted frame-by-frame estimates of the subject’s head position and heading (*θ*_Head_); the ground-truth azimuth of the target (*θ*_Target_); the subject-relative azimuth of the target (*θ*_Head-Target_); and crucially, the *angular (azimuthal) error* (*θ*_Err_) between the target center and the subject’s heading (Figure 1C). Data points from blurred or occluded frames were imputed using the modified Akima cubic Hermite interpolation. This yielded an azimuthal time course of head orientation, normalized to target azimuth, which provided the basis of further analyses (Figure 1D). See Appendix 2 for further detail.

#### Echoacoustic Click Extraction

To align the subject’s clicking and self-motion behavior, we extracted the audio track from each trial’s video recording and analyzed amplitudes within a 1001-sample (48-ms) sliding window. Each window was represented using mel-frequency cepstral coefficients (MFCCs) and classified using a binary support vector machine (SVM) manually trained to distinguish click events from background and environmental noise. This method enabled robust detection of clicks, characterized by short, high-energy transients, but distinct from equipment creaks or other incidental impact sounds. Detected click timestamps were then mapped to the head motion data to estimate the observer’s heading at the time of click emission. Finally, for click-acoustic analyses, we aligned head-centric in-ear recordings with those from the overhead camera via a cross-correlation procedure, then extracted 200ms of binaural audio centered at each aligned click timestamp. Head-centered click intensities were indexed by the peak dimensionless amplitudes of the raw recordings within these windows. See Appendix 3 for a more detailed description of the extraction model.

### Data analysis

#### Head Kinematics

For each trial, we measured the observer’s heading relative to the target (i.e. head angular error, *θ*_Err_) throughout the trial, as described above and in Figure 1. *θ*_Err_ in the final video frame of the trial, indicated by the button-triggered LED illumination, served as the participant response for that trial. The *angular error change*, *Δθ*_Err_, indexed the change in angular error magnitude between the start and end of a trial. Note that for results below, we report *θ*_Err_ as an absolute value. We measured time-varying aspects of the observer’s kinematic and active sampling behavior, including the number and rate of clicks produced and angular *heading reversals*, directional changes in head motion. These variables allowed us to more directly quantify the relationships between sensory sampling (clicks produced), convergence dynamics (head movements toward the target), and the time course of a trial. The reversals, taken as an index of goal-relevant movement, were computed by identifying local extrema in the head angular trajectory, separated by at least 0.5 s and with magnitude at least 2°. To formally characterize these relationships, we fitted an exponential decay function (Eq. 1), to the data in each condition, omitting the earliest 5% of indices to minimize distortion from single- or few-trial outliers. The function is of the form

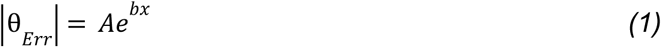

where absolute *θ*_Err_ is a function of *x*, given an offset parameter *A* and a decay constant *b*. In our case, *A* is the expected angular error at the end of the trial, when the time- or click-discretized countdown *x* has reached zero, at a rate governed by *b*. The relationship can be characterized with a “half-life” time constant as in Eq. 2:

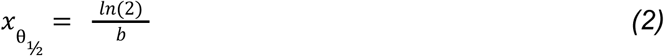

in which *x*_θ½_ is the time or clicks over which *θ*_Err_ decreases by half. We compared exponential vs. linear fit by comparing Akaike Information Criterion (AIC) for each model. (Burnham and Anderson 2002). Finally, to assess temporal dependencies between click timing and head reversals, we cross-correlated click and reversal timecourses in each trial. To do this, we resampled the video and audio data at 100 Hz (10 ms bins) to represent click and reversal events as sparse binary time series. Adapting methods used to compare neuronal spike trains (Perkel et al. 1967; Peyrache et al. 2015), we computed trial-wise cross-correlograms over lags of up to ±2 s, with coincidence counts at each lag summed across trials, then normalized against those expected by chance.

#### Click Directivity

To evaluate whether the observed behavior was consistent with an acoustic slope- or intensity-maximization strategy, we examined click heading histograms, both pooled and divided into quartiles of the trial time course. The goal was to estimate, based on these distributions and the acoustic directivity of the clicks, which part of the click profile was used to ensonify the target. We did not measure EB’s click directivity directly but used previously modeled click directivity functions (Thaler et al. 2017), with a peak at 0° and greatest slope roughly 30° off-axis. Our rationale was that a slope-maximizing localization strategy would most likely produce a bimodal distribution of click headings aimed ∼30° to either side of the target (Yovel et al. 2010), while an intensity-maximization strategy would result in click headings clustered unimodally around target center.

#### Statistical Significance

To avoid assumptions about normality or equality of variance between conditions, statistical within-subject effects of condition on response accuracy and other variables were tested via a distribution-free permutation-based ANOVA (“permANOVA”; e.g. Anderson 2001), which compares the observed sums of squares to a null distribution generated by shuffling group labels 10,000 times. Similarly, to test relationships between, e.g., within-trial click position and interclick interval or peak amplitude, we correlated click information pooled across trials and tested for significance via 95% confidence intervals (CIs) computed by bootstrapping the sample 10,000 times. For cross-correlations, significance across lags was estimated against a null distribution generated with a jitter permutation procedure (N=10,000) that randomized fine-scale click-reversal temporal relationships while preserving event count and coarse temporal structure within each time course. Each event was independently displaced by a uniform random offset within ±0.5 s (±50 samples), scrambling coupling finer than this window while retaining slower rate structure, including shared trial-onset locking. The test statistic was maximum absolute (z-scored) cross-correlation, controlled for family-wise error at α=0.05 (Maris and Oostenveld 2007; Nichols and Holmes 2002).

## RESULTS

*<u>Localization accuracy is gated by active echolocation and modulated by target size</u>* Figure 2A–C shows conditionwise angular head trajectories for each trial, and the error distributions of the responses. The average signed angular error did not vary by condition (1-way permANOVA, *F*_(2,178)_=1.17; *p*=0.314), suggesting responses were not systematically biased by our manipulations. However, precision varied widely, and the absolute angular error was significantly larger in both Small Target (24.56°) and No-Click (35.81°) conditions than in the Big Target condition (9.47°; *F*_(2,178)_=29.51; *p*=0.0001; both pairwise *p*=0.0003). Additionally, *Δθ*_Err_ varied significantly (F_(2,178)_=25.26; *p*=0.0001), with post-hoc tests confirming that angular error did not improve significantly in the No-Click condition, but did so in both clicking conditions, with a greater improvement for the Big Target (both *p*<0.05, Bonferroni-Holm corrected). Finally, EB did not trade speed for accuracy: Big Target trials lasted less than half as long as Small Target trials (9.3 s vs 23.2 s duration; 13.9 s difference; *p*=0.0003, Bonferroni-Holm corrected). In sum, as expected, EB echolocalized the large target more precisely and more quickly vs. the small target, and could not converge at all on the large target without active echolocation. We therefore focus subsequent analyses on the two active echolocation conditions.

**Figure 2.**
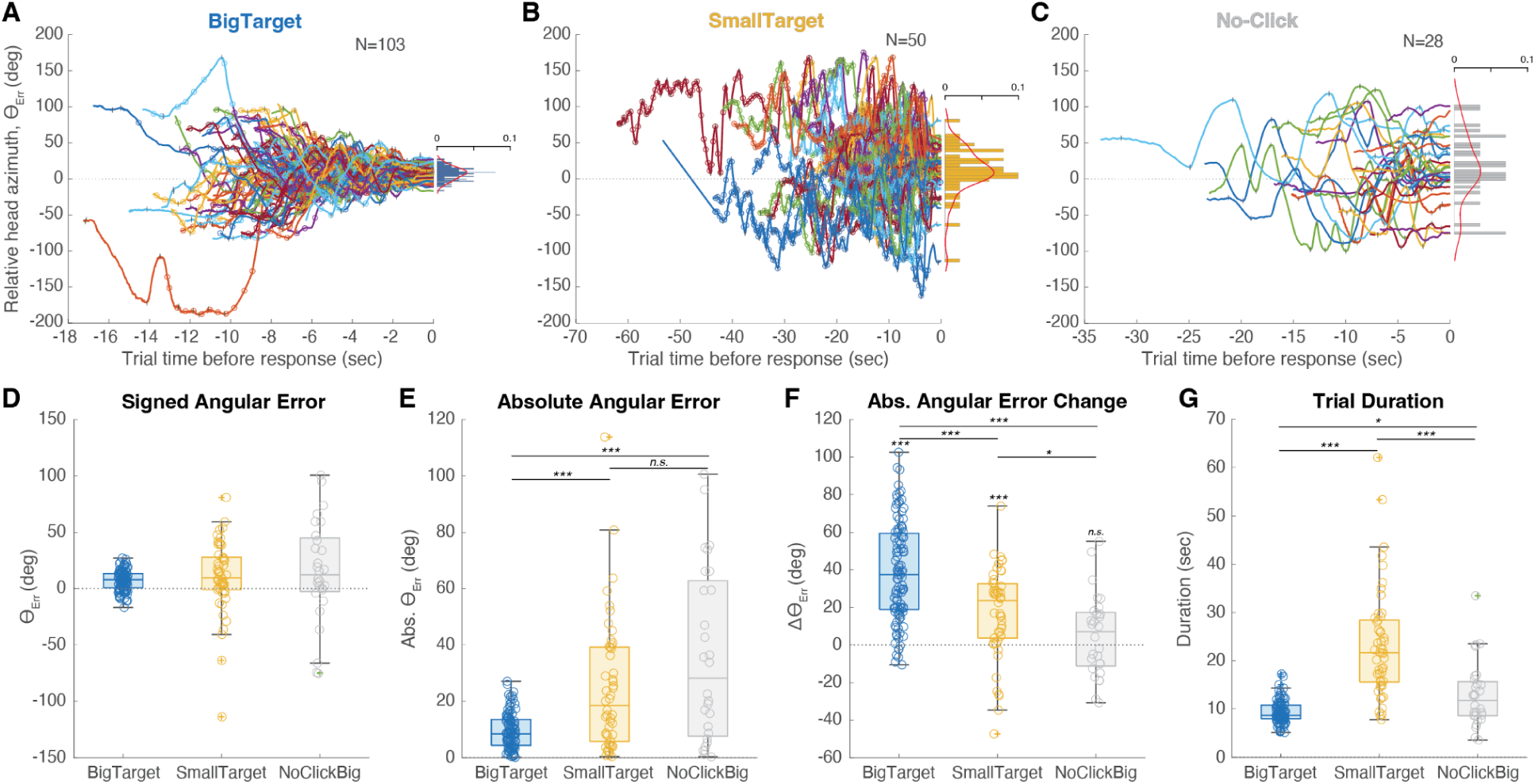
Angular heading errors *θ*_Err_ by trial and condition. **A-C**. Trialwise *θ*_Err_ trajectories by condition, annotated with clicks (‘o’) and heading reversals (‘|’). Response heading (final *θ*_Err_) histogram y-axes index normalized proportions. **D-E**. Signed and absolute *θ*_Err_ by condition. Box-and-whisker plots indicate median, IQR, and outliers (‘+’), overlaid with individual trials. Statistical comparisons indicate pairwise post-hoc tests to permutational ANOVA. **F**. Absolute *Δθ*_Err_ from first to last trial timepoint (positive value denotes improvement, i.e. reduced *θ*_Err_). Boxplot statistics indicate 1-sample tests against zero. **G**. Trial durations from first to last trial timepoint. For all analyses, significance determined by permutation, Bonferroni-Holm corrected. * = p<0.05; ** = p<0.01; *** = p<0.001.

### Target size modulates clicking behavior, but not head reversals

Active-sensing behavior, evaluated via kinematic and click analyses shown in Figure 3, indicates that EB produced fewer average clicks per trial (12.7 vs 32.6) and at a slightly slower rate (1.37 vs 1.81 Hz) for Big Target vs. Small Target conditions, respectively (both *p*=0.0003, Bonferroni-Holm corrected). By contrast, while the number of heading reversals — indexing EB’s side-to-side head motion — per trial differed systematically by condition, reversal *rates* did not differ between Big Target and Small Target conditions (0.87 and 0.86 Hz), though both were slightly faster than the No-Click condition (0.50 Hz, *p*=0.0003).

**Figure 3.**
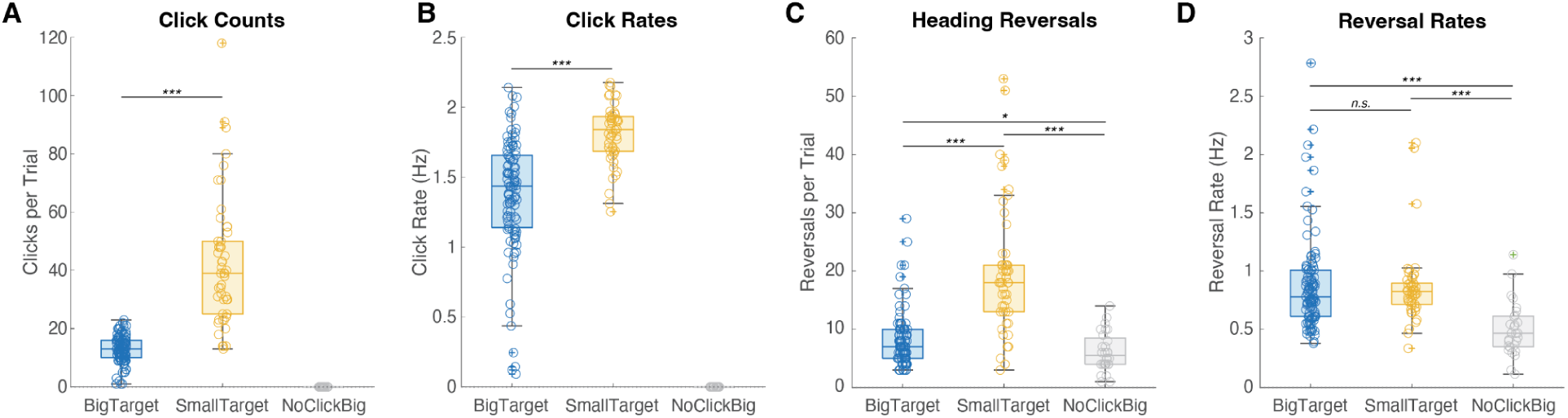
Echolocation click and head-movement dynamics. **A**. Trialwise click counts were higher and more variable in the Small Target vs. Big Target condition, and click rates (**B**) were significantly but only moderately higher, suggesting a sampling rate typically falling between 1–2 Hz. **C**. EB made more heading reversals in Small Target vs. the other two conditions, but accounting for trial duration (**D)** shows the average reversal rates were equal in the two active conditions, and lower in the No-Click condition. * = p<0.05; ** = p<0.01; *** = p<0.001.

Within-trial clicking and head movement dynamics corroborate this pattern when ordering data by click position index. Inter-click intervals (ICIs, the time elapsed since the previous click) remained constant until reliably rising in the last 3–4 clicks of a trial, as indicated by the click index at which the ICI-index correlation reached significance (Figure 4A). Inter-reversal intervals and click amplitudes did not vary systematically within a trial (Figure 4B,C), though click amplitudes overall were higher in the Small Target condition (p=0.0001). Finally, for each target condition, the trialwise click-reversal cross-correlation function, normalized to z-units, varied between about -2 and 3 for lags up to ±2 s. A few spikes approached the significance threshold at ɑ=0.05, especially in Small Target trials (Figure 4D), but overall the cross-correlogram fell short of the bounds of the surrogate null distribution (p_Global_ = 0.387 and 0.051 for Big and Small Targets, respectively). In sum, these results indicate that click rates varied from ∼1–2 Hz between conditions and trials; that they were constant *within* trials until a few clicks before the response; that click intensity varied across conditions but not within trial; and that head reversal kinematics were invariant both within trials and across conditions, and largely independent of individual click timing.

**Figure 4.**
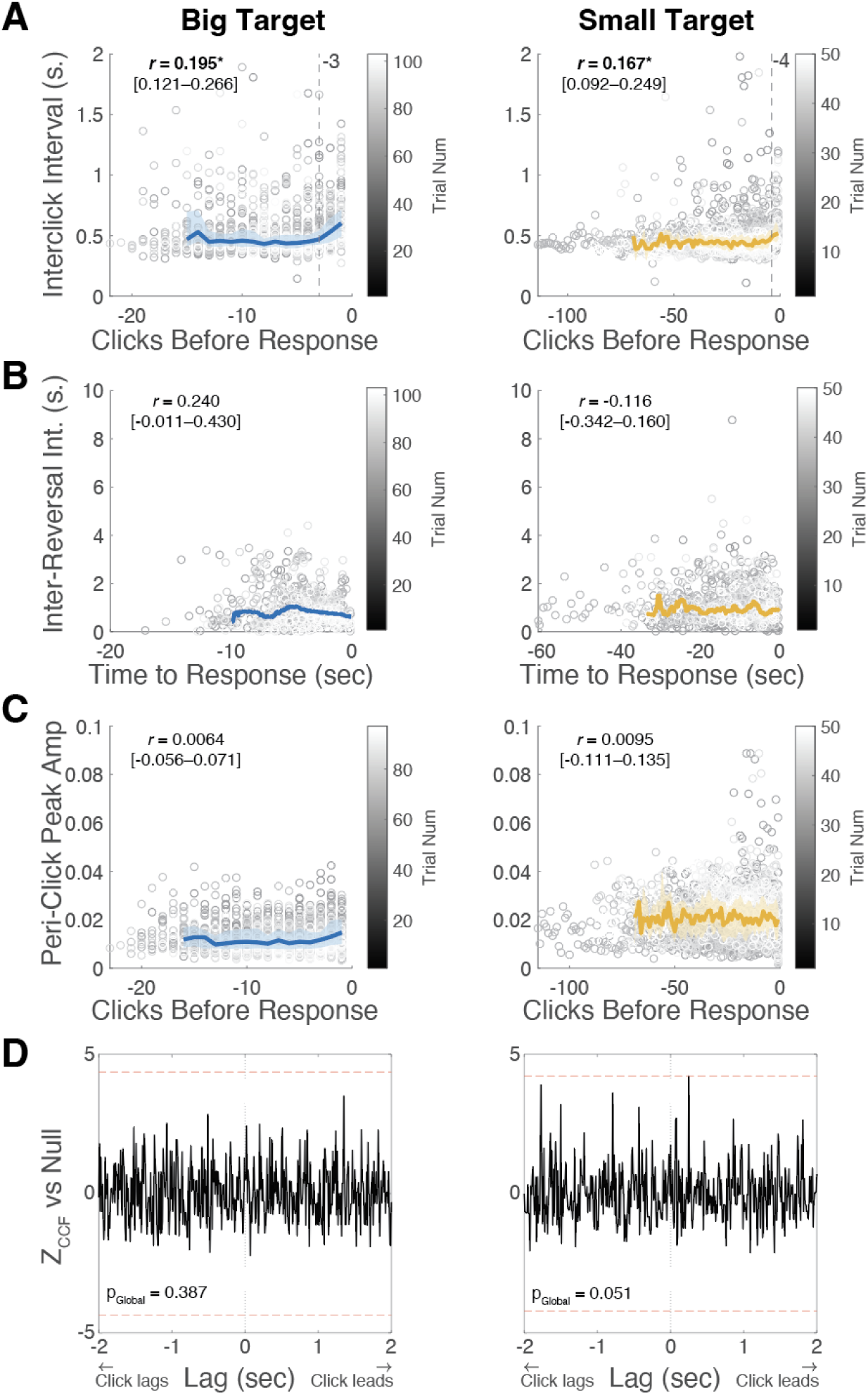
Within-trial click and kinematic dynamics, pooled by condition. **A.** ICIs (interclick intervals, the time elapsed since previous click) as a function of click index prior to response. Vertical dotted line: correlation significance breakpoint. **B**. IRIs (inter-reversal intervals, time between head direction changes) as a function of time to response. **C**. Peak amplitude of click audio, raw units. Colored plots in all panels: median values by click index or moving average across a 2-s sliding window. Shaded patches: Interquartile ranges by click index. Grayscale gradient: trial index. Within-trial Pearson correlations include bootstrapped 95% confidence intervals in brackets. To reduce distortion from few-trial outliers, the earliest 5% of data points are omitted from analyses. **D**. Click-Reversal cross-correlation function (CCF) z-scores vs. a jitter-permutation null distribution within a ±2 s lag window. Red dashed lines indicate pooled significance thresholds. p_Global_ = p-value of significance anywhere in the window. See Methods for details.

### Azimuthal convergence dynamics are consistent with exponential decay

Figure 5 shows absolute *θ*_Err_ plotted against heading reversal timestamps and click indices, aligned to the trial-ending response. The *θ*_Err_ progression in each plot is shown averaged over a 0.5-s sliding window for reversals and averaged by each index position for clicks. Additionally, the exponential decay fit (Eq. 1) is shown overlaid in red. Although EB’s oscillating head motion produced widely varying head positions, the fit closely matches the average convergence.

**Figure 5.**
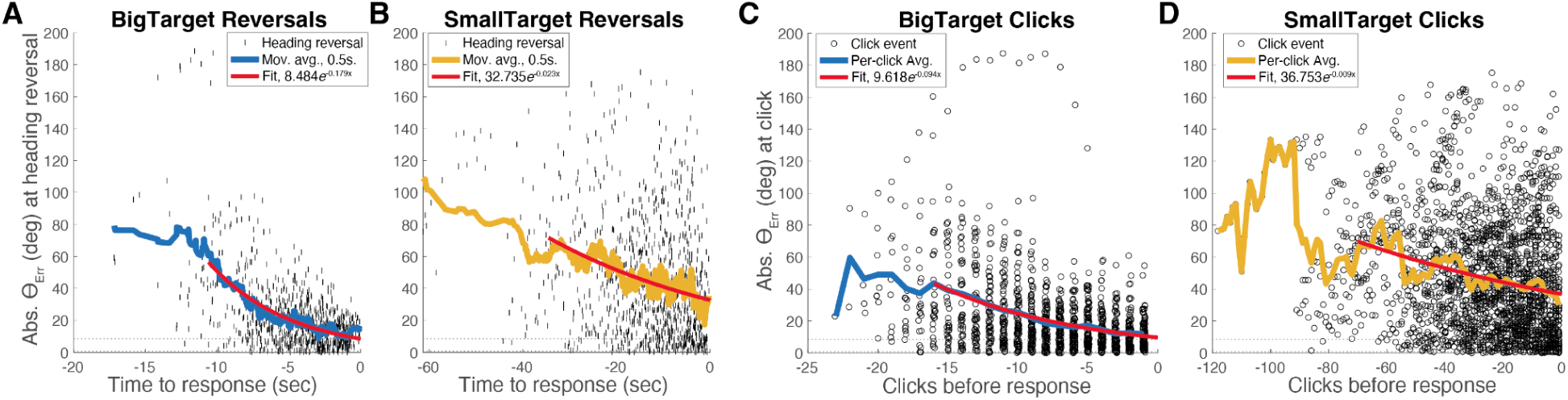
Convergence dynamics from pooled trials within clicking conditions, showing |*θ*_Err_| at heading reversals vs. time to response (**A**-**B**) or at click emission vs. clicks to response (**C**-**D**). Thick color-coded lines are moving averages in a 0.5 s sliding window. Horizontal dotted lines indicate azimuthal half-angles subtended by large (±8.5°) and small (±0.7°) targets. Red traces are model fits suggesting |*θ*_Err_| at reversals halving every 3.9 s and 30.1 s, or at clicks every 7.4 clicks and 77 clicks for large and small targets, respectively.

Comparing AIC, using a convention in which positive *Δ*AIC favors an exponential vs. linear model (see Table 2), yielded strong support for an exponential profile for large targets (*Δ*AIC = 40 and 37.3 for click-number-indexed and reversal-time-indexed convergence, respectively). For small-target trials, click-number-indexed convergence weakly favored an exponential model (*Δ*AIC = 2.1), while reversal-time-indexed convergence was not distinguishable from a linear profile. A control analysis of no-click reversals showed no convergence and, as expected, no favored functional form (*Δ*AIC = -0.2).

**Table 2.**
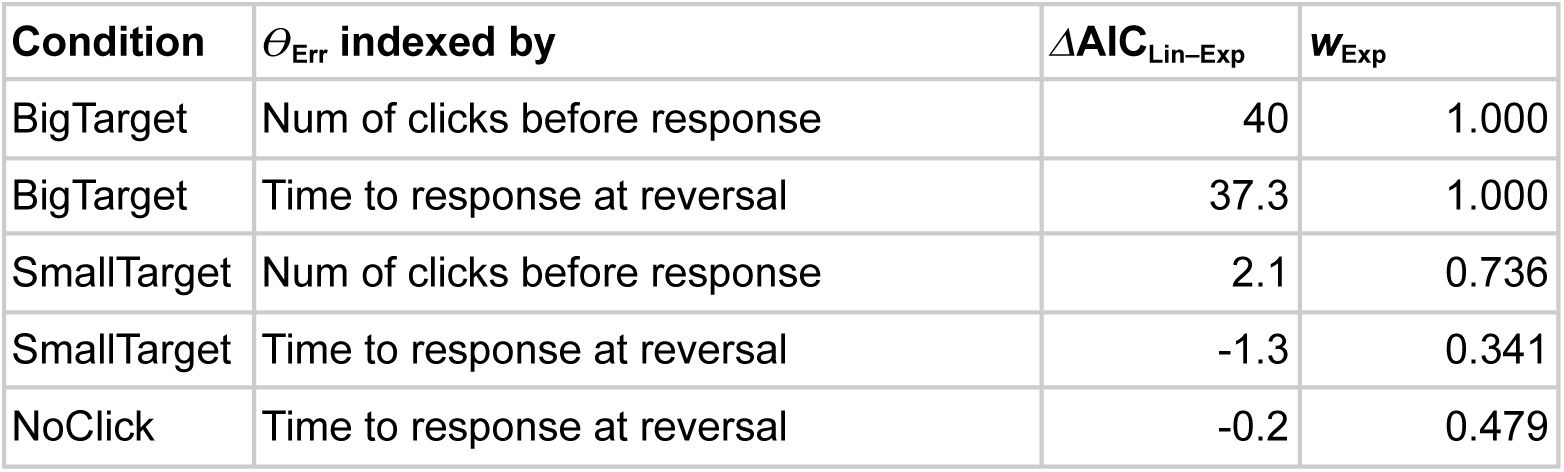
Model comparison of AIC for exponential fits plotted in Figure 5 vs linear fit.

Angular “half-lives” (Eq. 2) of reversal- and click- indexed convergence were estimated, respectively at 3.9 s and 7.4 clicks for Big Target trials, and 30.1 s and 77 clicks for Small Target trials. These analyses suggest that given a sufficiently strong echo, EB approached the target faster earlier in a trial than later, but that the proportional rate of convergence was constant, and well predicted by an exponential function relating it to clicks. For both the reversal and click analyses, the time or clicks to halve *θ*_Err_ increased by about an order of magnitude for small vs. large target trials.

### Unimodal click distribution suggests maximum-intensity ensonification strategy

How did EB deploy the sound energy of echolocation clicks to localize the target? To estimate the echoacoustics of ensonification, we compared the relative angle of each click (*θ*_Err_ at the time of emission) with previous estimates of click directivity derived from modeling many click recordings. In Figure 6A, click heading polar histograms, color-coded by target condition, are overlaid with a representative echolocation click amplitude directivity plot (solid black line, “EE1” model parameters (Thaler et al. 2017)). The two-lobed dotted plot represents the instantaneous slope of the function, maximal near ±30° relative to center. Both Big and Small Target histograms were unimodal and peaked closer to the center than either eccentric locus (Figure 6A,B), consistent with maximum click intensity directed toward the target. Next, we separated click headings into quartiled sub-epochs between trial start and finish. The similarly unimodal distributions (Figure 6C) suggest this ensonification strategy was constant throughout the trial.

**Figure 6.**
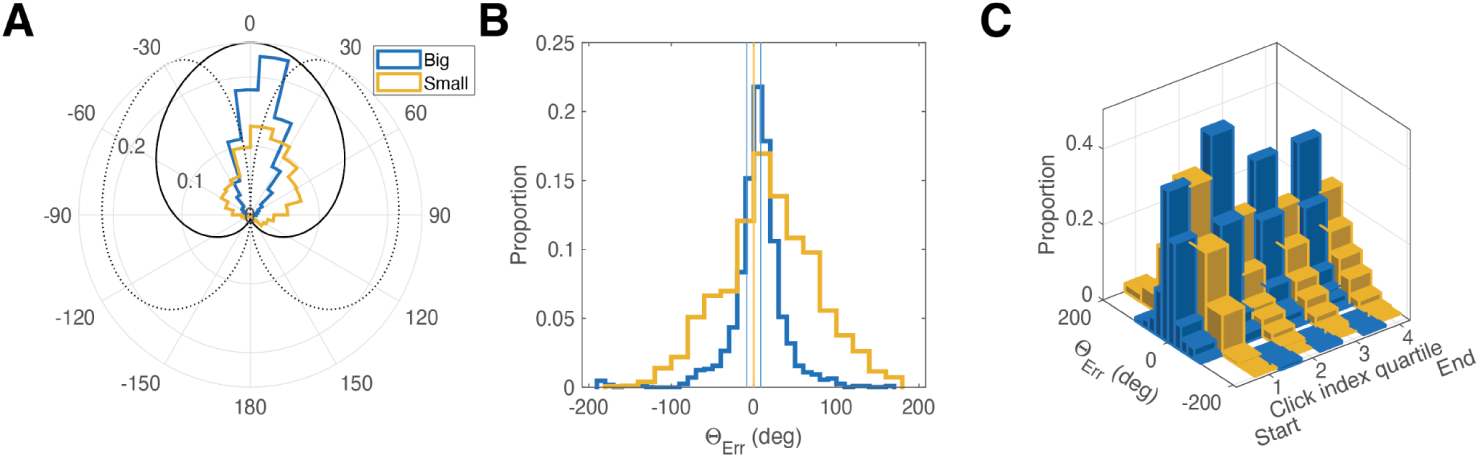
Click headings relative to targets by condition. **A.** Polar histogram of click headings relative to Big (blue) and Small (yellow) targets. Radial axis: proportion of total clicks. Solid black trace: representative human click intensity directivity (Thaler et al. 2017). Dotted trace: directivity slope (instantaneous derivative). **B.** Histogram of all clicks by condition; color codes same as (A). Vertical traces represent edges of targets relative to center. **C.** Click heading distributions over trial, quartiled from 1 (early in trial) to 4 (last quartile before response).

## DISCUSSION

In this study we characterized, for the first time, the active-sensing dynamics of echoacoustic target acquisition and localization in a proficient blind human echolocator. We found faster and more precise localization for large vs. small targets, and no convergence at all when clicks were restricted, indicating that head-orienting precision was sensitive to both target and sensor manipulations. Echolocation clicks were more numerous and intense for small-target trials. At the same time, within trials, head movements and clicks were produced at constant but independent (or weakly dependent) rates, with click emissions slowing slightly in the final few clicks prior to the response. Head movements were roughly oscillatory, but converged toward the target along an exponential trajectory, most clearly evident for the Big Target condition.

Finally, click headings indicated direct head-on ensonification, an approach consistent with an echo intensity-maximization, rather than, e.g. slope-maximization strategy.

### Adaptive and invariant active-sensing dynamics

The head kinematics and click sampling we observed reveal an intriguing dissociation between adaptive and invariant aspects of the sensorimotor loop. For both target sizes, EB reversed head direction at a constant rate, and produced clicks at a constant (albeit condition-dependent) amplitude and rate, until the final 3–4 clicks of a trial. That is, EB converged reliably toward the target using sampling behavior modulated by target size but largely invariant within a trial. Additionally, clicks did not seem to directly drive head reversals: the cross-correlations did not reach family-wise significance in either condition, though the Small Target approached it in isolated time bins within the ±2 s lag window (most closely at a click-lead offset of ∼0.25 s). The near-threshold spikes are thus unlikely to reflect a tight click-to-reversal coupling.

In a classic early echolocation study with a similar experimental setup to ours (Rice 1969), participants made an unspecified number of clicks while stationary and then silently turned their heads toward the perceived target azimuth. Much more recently, a series of virtualized experiments demonstrated that free head movements aided echoacoustic orientation to align with a hallway (Wallmeier and Wiegrebe 2014). Drawing from prior results such as these, a simple click-driven sequential triangulation model of echoacoustic orientation might predict head reversals to be elicited (and therefore disproportionately preceded) by clicks. Instead, both were distributed evenly throughout each trial, such that a given click did not, by itself, predict the observer’s heading adjustments or the imminence of a final response. Still, clicking behavior was clearly necessary to successful performance overall. Our interpretation does not require strict independence, but our results are consistent with an account in which a multi-click integrative process gradually reduces angular error.

### Exponential convergence profile and click-wise evidence accumulation

The structure of EB’s heading convergence pattern has several implications. First, as seen in Figure 5, the exponential decay model closely captures the *moving average* of head orientations across trials rather than the instantaneous, widely varying azimuths (at click emissions or heading reversals) to which it was actually fitted. The model may thus capture echoacoustically driven convergence dynamics dissociable from a particular head-movement strategy used by EB. Second, the time constant of angular error reduction — a “half-life” of ∼7.4 clicks for Big Target trials and ∼77 clicks for Small Target trials — suggests an estimate for the informational value of clicks that is broadly modulated by task conditions but follows a constant rule within a trial. Specifically, in this case, converging toward the smaller target required over 10 times as many click samples as an equivalent error reduction required for the larger target. The weak support for exponential vs. linear convergence in the Small Target condition is an unsurprising corollary, as EB traversed less than a single angular half-life (by either measure) in those trials. Third, this pattern contrasts with linear click-wise improvement in left-right lateralization thresholds reported in a headphone-based psychophysical experiment (García-Lázaro and Teng 2026). In a head-fixed paradigm, using synthetic echoacoustic stimuli, the information in each click per trial was identical. By contrast, in the present study, as a free-moving head reduces its angular error, successive clicks ensonify the target at (and echoes return from) a progressively smaller azimuth, which reduces the binaural timing and level differences from which an angular correction can be planned. The experimental setup and resultant echoacoustics thus predict the proportionality of *θ*_Err_ to its rate of change, a relationship found in other sensorimotor refinement scenarios such as visual reaching (Burge et al. 2008), saccadic gain adaptation (Straube et al. 1997), or speech pitch correction (Parrell et al. 2019).

### An intensity-maximization ensonification strategy?

What part of the echolocation signal is most useful to the observer? In two prior examples of off-axis echolocalization (Thaler et al. 2022; Yovel et al. 2010), the relevant factor was the rate of change in the signal, a well known sensory principle even at the level of neuronal tuning curves (Verghese et al. 2012; Butts and Goldman 2006). One consequent prediction was that “echolocators using clicks in a scenario in which they can move their heads may orient their heads slightly away from the target when their goal is to localize the target’s position” (Thaler et al. 2022). Based on the distributions of headings (and thus peak click intensity) in Figure 6, our results are more consistent with an intensity- rather than a slope-maximization strategy. The task instruction to point the head at target center may have artificially constrained head movements, but the quartiled click index histograms (Figure 6C) argue against this: in all portions of the trial, the click heading distribution was unimodal and centered on the target, never taking on the bimodal distribution characteristic of a slope-maximization strategy (Yovel et al. 2010). Nevertheless, our results do not necessarily contradict the utility of signal slope.

Indeed, under suboptimal conditions, the echolocating bats in (Yovel et al. 2010) shifted to ensonifying their navigational targets with the peak beam intensity rather than slope, a dynamic tradeoff driven by available signal-to-noise ratio. In the present study, EB may have combined the greater SNR of on-axis ensonification with the freedom to point that axis at different parts of the target. Speculatively, a head-free intensity-maximizing strategy may still rely on changes in echo signal, but indexed across proprioceptive cues (i.e. head position changes), rather than azimuthal eccentricity. This interpretation would be consistent with the side-to-side exploratory movements EB exhibited and progressively refined in each trial.

### Summary, limitations, and future work

The term *echolocation* is not monolithic, but rather comprises a family of behaviors and percepts that are observer- and task-dependent. This work represents a step toward characterizing not just observable strategies, but candidate acoustic and perceptual mechanisms mediating this behavior. Taken together, the head scanning, click production, and convergence dynamics suggest a closed-loop coarse-to-fine tuning process driven by the integration of evidence from repeated sensory samples, with behavioral adjustments gradual rather than tightly coupled to individual clicks. Aside from task-related convergence, within-trial dynamics are stable: click intensity and rate vary by trial and task difficulty, but not systematically within a trial. Late-trial slowing is possibly tied to response preparation. Head movements oscillate at a fixed rate, aiming the highest-intensity portion of click energy toward the target.

Importantly, in focusing on an early-blind proficient echolocator, we do not claim to definitively characterize how echolocators behave as a population. Rather, this study explores a representative case of a process whose dynamics are not well understood. Notably, a self-reported blind echolocator did not employ these sensorimotor dynamics successfully in our task (Appendix 1), implying the necessity of expertise beyond casual practice or blindness per se. In this vein, testing additional proficient and novice participants would allow a fuller characterization of the ways in which echolocation expertise manifests under different conditions and in different stages of training. Further studies may also clarify how precisely echoacoustic and proprioceptive cues combine to represent allocentric vs. egocentric target location. The resolution of our present data cannot directly address this question, but it is intriguing in light of evidence that free-flying echolocating bats predict trajectories of moving targets rather than relying on the most recent echo (Salles et al. 2020). Questions such as these are ripe for another fertile research direction — the quantitative modeling of the sequential dynamics characterized here, e.g. with a Bayesian or other recurrent approach (Krasovskaya et al. 2025; Yang et al. 2016; Renninger et al. 2007). This would establish a formal framework for the active-sensing architecture, facilitate the identification of behaviorally salient parameters, and distinguish, e.g., fundamental dynamics from idiosyncratic strategies. Finally, a practical implication of this work is that echolocation learning in orientation and mobility contexts may benefit from integrating head-scanning strategies with training in echoacoustic perception.

## ACKNOWLEDGMENTS

We thank Joshua Miele and James M. Coughlan for valuable comments and discussion in the conception and development of this project, and our participants for their time. GF’s contributions were made during his affiliation with SKERI. This work was supported by the Foundation for Ophthalmology Research and Education International (ST); NIH Training Grant 5T32EY025201-03 (SKERI/ST), NIH Grant 1R21EY032282-01 (ST), and RERC grant 90RE5024-01-00 (SKERI).

## APPENDIX 1 SUPPLEMENTAL DATA

**Figure A1.**
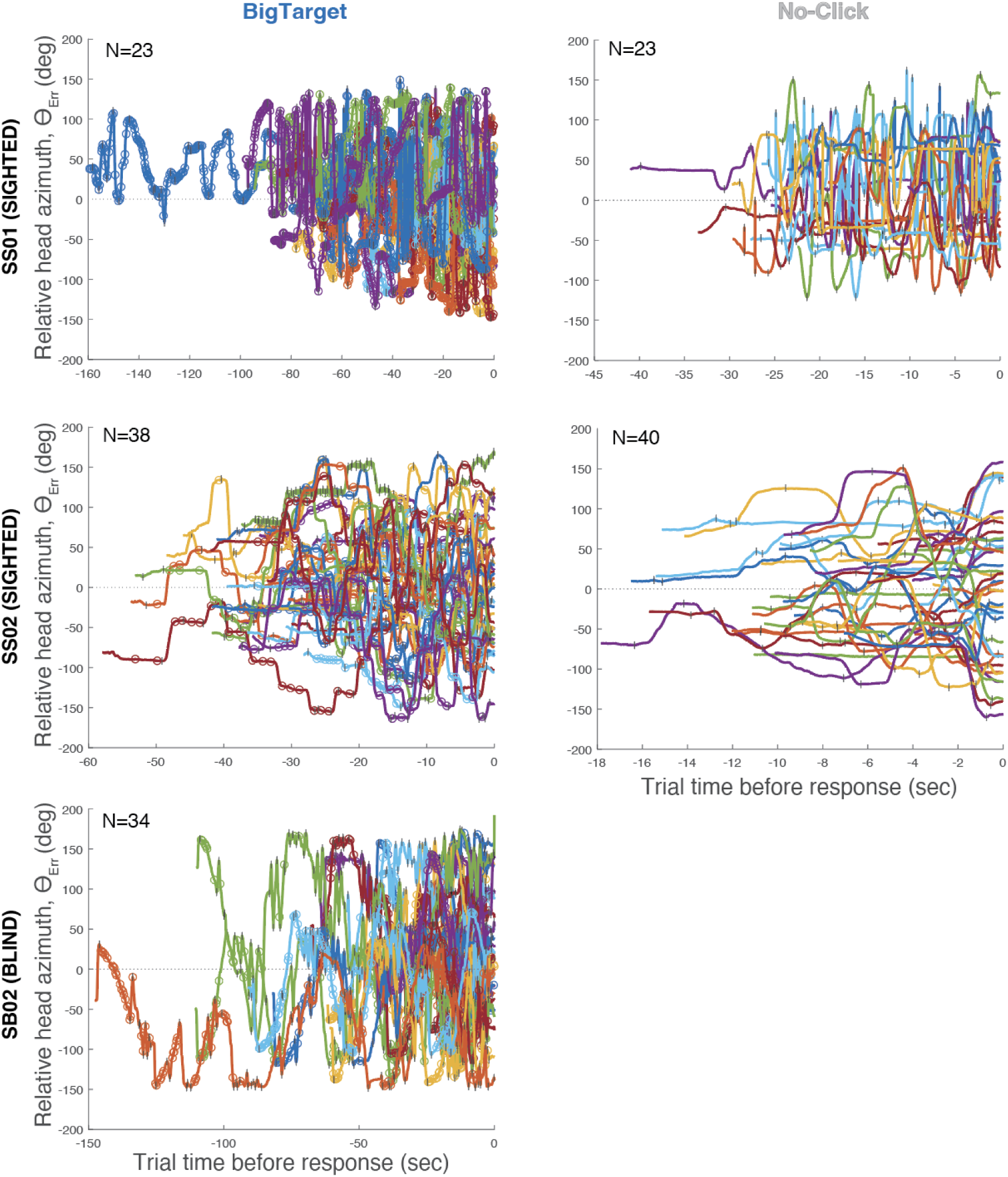
Trialwise angular error trajectories for two additional sighted (SS01, SS02) and one blind (SB02) participant. Plotting conventions as shown in Fig. 2 of main text. Panels show trial counts by condition.

**Figure A2.**
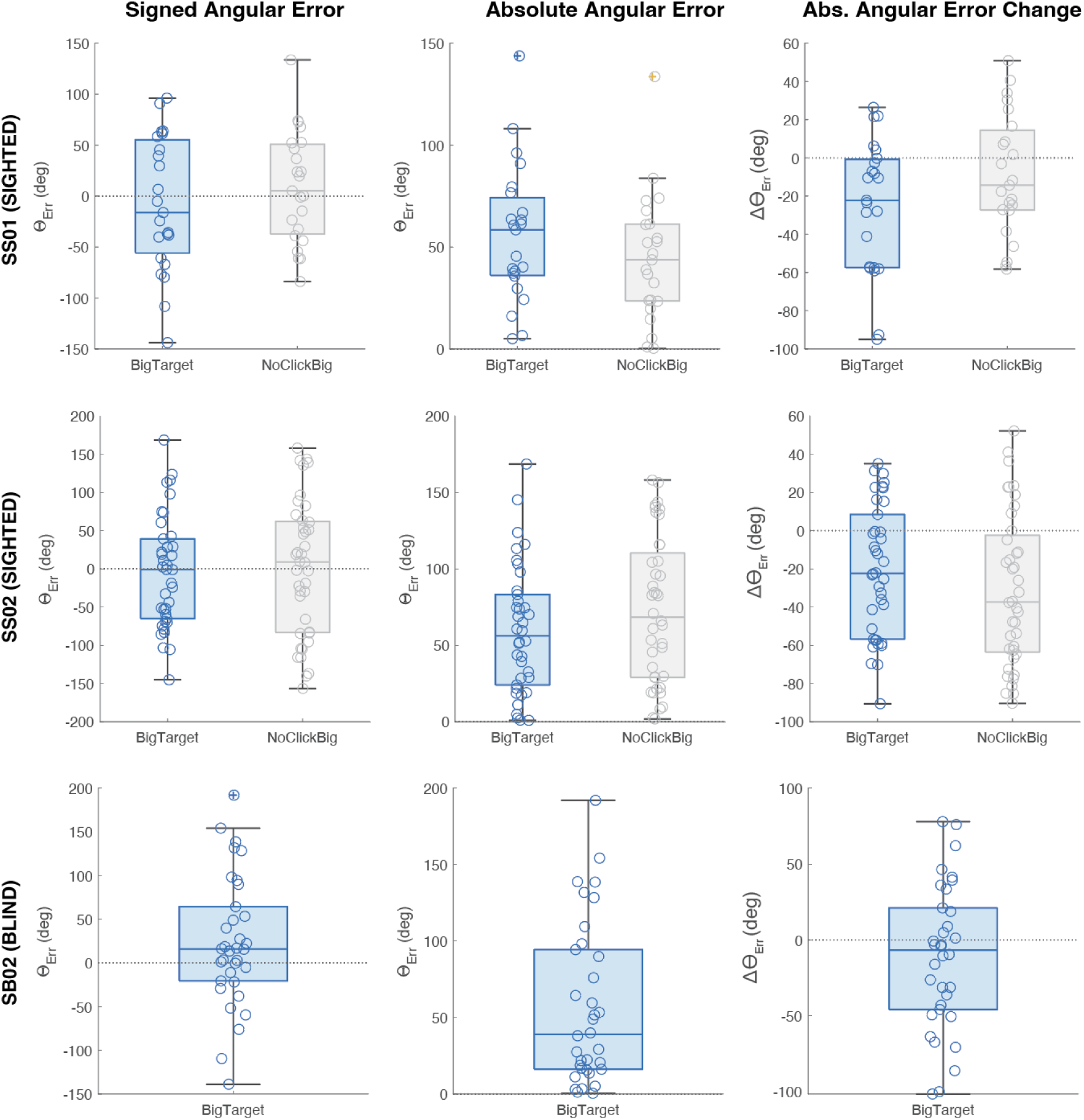
Behavioral accuracy for two additional sighted and one blind participant. Plotting as shown in Fig. 2 of main text. No improvement was observed in any of the tested conditions. Negative *Δθ*_Err_ for SS01 and SS02 (larger ending vs. starting *θ*_Err_) may reflect a narrow starting distribution.

## APPENDIX 2 HEAD AND TARGET TRACKING PIPELINE

### Overview of the Approach

Our video-based tracking system was developed to facilitate the automated analysis of human echolocation experiments in which participants employ self-generated tongue clicks and subsequently interpret the returning acoustic echoes to determine the angular direction of a reflective target. The principal objective is to quantitatively evaluate the precision with which participants estimate the angular position of the target via head-pointing at the moment they signal completion of the echolocalization task, as indicated by a button press.

### Experimental setup

A calibrated GoPro camera (HERO5, GoPro Inc., San Mateo, CA) is mounted above the participant’s chair, aligned parallel to the ground. Thus its field of view (FOV) provides a top-down view of both the participant and the target apparatus, but not the target itself, which is outside the camera FOV.

The seated participant wears a head-mounted ArUco marker, enabling precise tracking of head position and orientation throughout the session. The echo-reflective target consists of a flat rectangular panel mounted on a rotating rod. The camera, rod rotational axis, and participant are all roughly collinear. Thus, the rod’s rotation changes the azimuth angle of the target between trials, while its mean distance from the participant remains fixed at ∼1 m. A small red-taped element is attached to the rod in such a way that it is 1) parallel to the rod and 2) always in the camera’s FOV. Combined with the known geometry of the setup, this visual cue thus provides a reference for extracting target azimuth in image coordinates from any given video frame.

At the end of each echolocation trial, the participant presses a button that activates an LED mounted within a fixed corner of the camera FOV. The LED illumination marks the moment when the participant believes they have localized the target, and thus determines the end of the trial. The head orientation during this frame is taken as the localization “response” from which angular errors and other parameters are computed. Subsequent missing marker detections for several consecutive frames indicate a transition to a new trial.

### Center of rotation estimation

The system first determines the rotation center of the rod by tracking a red-taped marker across image frames and fitting an ellipse to its observed trajectory. The center of the fitted ellipse defines the geometric origin used for subsequent angular calculations. This calibration step is performed once during system initialization and reused for all trials in a given recording.

To obtain the marker trajectory, the rod is manually rotated while the camera records the motion. The red-taped marker is segmented in each frame using hue-based thresholding in the HLS color space, which provides robustness to illumination variations. For each segmented frame, the centroid of the marker is computed using standard blob geometry. The resulting sequence of centroid positions forms a closed path corresponding to the rod’s rotation, from which the ellipse parameters and rotation center are estimated.

### Detection and angle estimation

For each frame of the video, the system extracts the participant’s head orientation, the target direction, and their relative angular relationship within a common image coordinate frame.

The original GoPro image is first undistorted using the calibrated intrinsic and distortion parameters. The red-taped guide attached to the rotating rod is segmented from the camera frame, and its position is used to infer the orientation of the rod in the image. Knowing the rod’s physical length and the camera’s pixel-to-meter scale (camera intrinsics), the system projects a line from the center of rotation of the apparatus *c* (previously determined during calibration) through the detected marker to obtain the target point *t*. The angle between this line and the horizontal axis defines the scene-level target azimuth

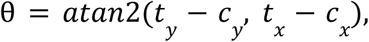

which represents the direction of the reflective panel relative to the apparatus’ pivot. The head pose is estimated independently from the head-mounted ArUco marker. Two key points are extracted: the head center *h* and a nose proxy point *n* indicating the forward-facing direction of the head. Head orientation in the image plane is given by the yaw angle

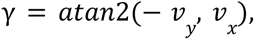

where *v* = *n* − *h* is the head-forward vector (the negative sign compensates for the downward orientation of the image *y*-axis). The head-centric target direction is defined by the vector *v*_*ht*_ = *t* − *h*, connecting the head center to the target position. Its angular direction,

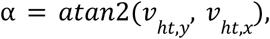

represents where the target lies relative to the participant’s current head position and serves as the reference for the perceived target direction. From these quantities, the system computes the instantaneous angular offset

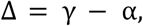

which quantifies how closely the participant’s head is oriented toward the reflective target at any given moment. (This quantity is referred to as the angular error in the main text.) The same geometry also allows the physical azimuth θ of the target to be expressed in the image coordinates. At the end of each trial the current values of α, γ, θ, and Δ are recorded to capture the participant’s final perceived alignment with the reflective target.

## APPENDIX 3 ECHOACOUSTIC CLICK DETECTION AND CLASSIFICATION

Audio segments centered around candidate click events were represented using Mel-Frequency Cepstral Coefficients (MFCCs), a widely used audio feature representation that captures perceptually relevant spectral characteristics while remaining robust to background noise and amplitude variations. MFCC features were extracted using the melfcc function from the Rastamat toolbox with its default configuration. This configuration computes a mel-scaled spectral representation of each audio segment using short-time Fourier analysis, applies a mel filter bank, and derives cepstral coefficients describing the spectral envelope on a frame-by-frame basis. The resulting MFCC coefficient matrix was flattened into a single fixed-length feature vector per audio segment.

**Figure A3.**
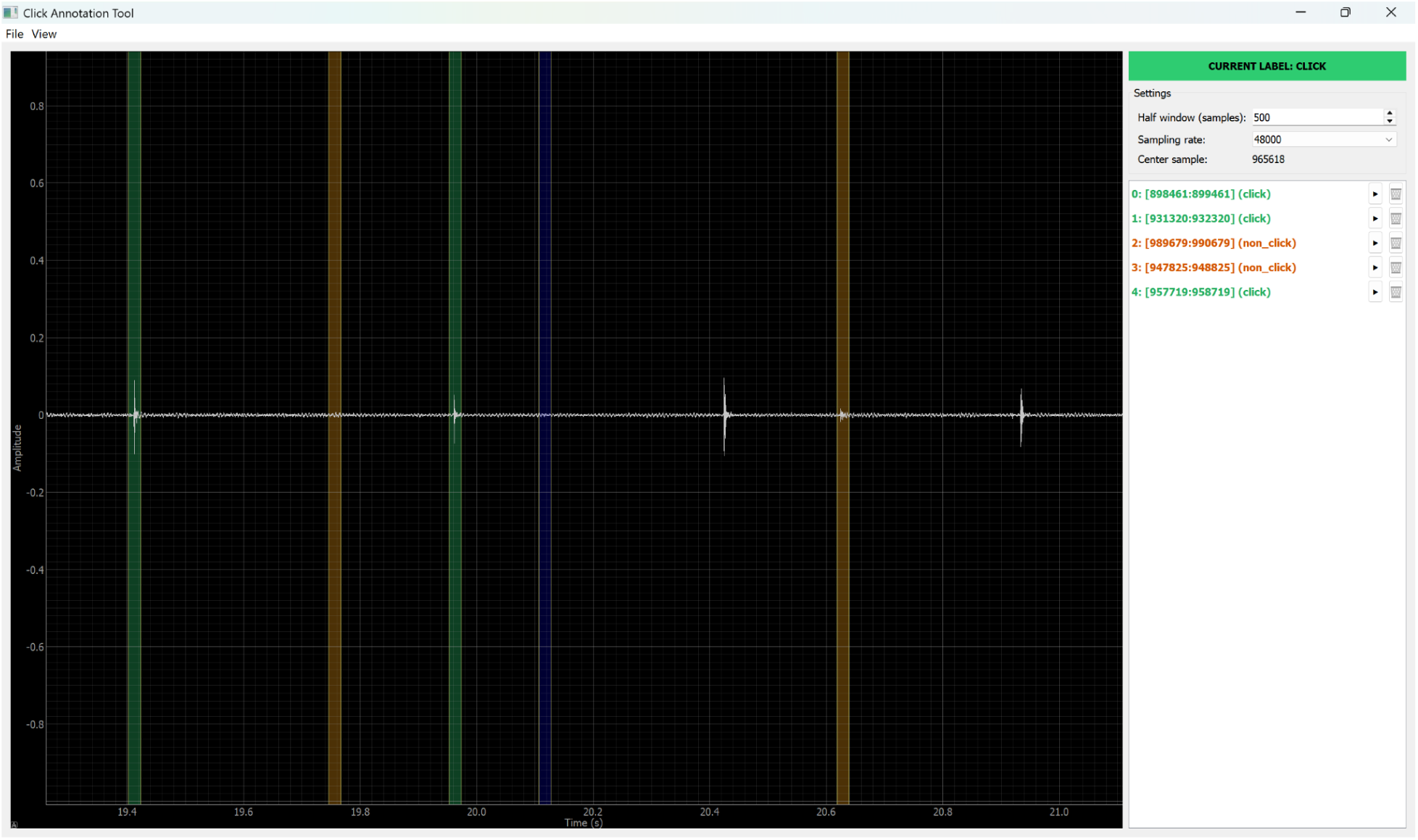
A screenshot of the audio annotation tool used to create the Click/NonClick dataset.

Click detection was formulated as a binary classification problem distinguishing click events from background and environmental noise. A Support Vector Machine (SVM) classifier was trained using manually annotated positive (click) and negative (non-click) samples. SVMs were selected due to their effectiveness in high-dimensional feature spaces and their ability to generalize well in regimes with limited training data, which is typical for manually annotated acoustic events. To avoid manual tuning and improve robustness, SVM hyperparameters were selected automatically using Bayesian optimization as implemented in MATLAB’s fitcsvm function. The optimization procedure iteratively evaluated candidate hyperparameter configurations using 10-fold cross-validation and selected new candidates based on an expected-improvement-plus acquisition function. This strategy balances exploration of the hyperparameter space with exploitation of promising regions, while explicitly accounting for uncertainty in performance estimates. The optimization was performed over a fixed evaluation budget, and the resulting model was selected based on cross-validated classification performance. The optimized SVM classifier replaces the previous amplitude-based heuristic and enables more robust detection of click events characterized by short, high-energy transients, while remaining compatible with the existing annotation and temporal alignment pipeline.

